# Simplified and refined murine model of reversible aortic constriction for characterizing cardiac functional recovery

**DOI:** 10.1101/2025.04.05.646758

**Authors:** Elnaz Ghajar-Rahimi, Pierre Sicard, Laura Castel, Craig J. Goergen, Thomas Moore-Morris

## Abstract

**Background:** Heart failure with reduced ejection fraction (HFrEF) is a major clinical issue with a poor prognosis. Structural and functional recovery, known as “reverse remodeling (RR),” has been observed in subsets of patients who respond favorably to treatment. However, progress in understanding the mechanisms behind RR has been hindered by a lack of animal models that accurately replicate this complex process.

**Objectives:** We introduced a convenient and reproducible modification to the widely used transaortic constriction (TAC) procedure that allows for refined, timed, and noninvasive aortic de-banding (de-TAC), and functional recovery.

**Methods:** A modified and releasable knot was used for an initial aortic constriction. Suture ends were left accessible from underneath the skin layer following the TAC surgery. After three weeks, deTAC was performed by re-opening in the skin layer and pulling on the suture to release the constriction. We assessed cardiac function, strain, and geometry with 2D and 4D ultrasound at baseline, 3 weeks after TAC, 1 week- and 4 weeks after deTAC. Tissue remodeling and cellular infiltration was characterized by quantifying fibrosis and fibroblast numbers via immunofluorescence.

**Results:** Ultrasound revealed that aortic de-banding was associated with both functional and mechanical recovery. Cardiac function and strain decreased due to pressure overload, exhibiting characteristics of systolic dysfunction, but recovered to near baseline values by 1 week after deTAC. However, left ventricular mass, fibrosis and fibroblast numbers remained elevated, despite functional recovery.

**Conclusion:** This simplified and refined TAC-based model of RR significantly facilitates and enhances research on cardiac remodeling and recovery.

**CONDENSED ABSTRACT:** A subset of heart failure patients (HF) respond favorably to treatment, displaying functional recovery via a process known as reverse remodeling (RR). Developing RR models will be key to characterizing currently elusive mechanisms driving this process and ultimately identifying potential therapeutic targets. Transverse aortic constriction (TAC) in rodents increases the afterload on the LV and is a common model of heart failure due to pressure overload. Here, we report a convenient and reproducible method that allows for precisely timed and surgery-free removal of aortic constriction (de-TAC). When performed 3-weeks after TAC, de-TAC produced significant cardiac functional recovery, with normalization of ejection fraction and strain assessed by 2D and 4D echocardiography, respectively. However, signs of adverse myocardial remodeling persisted, including hypertrophy, fibrosis, and elevated fibroblast numbers. In the relatively young field of cardiac RR, our method could facilitate investigations of functional, structural, cellular, and molecular changes underlying recovery from HF.

**VISUAL ABSTRACT:** 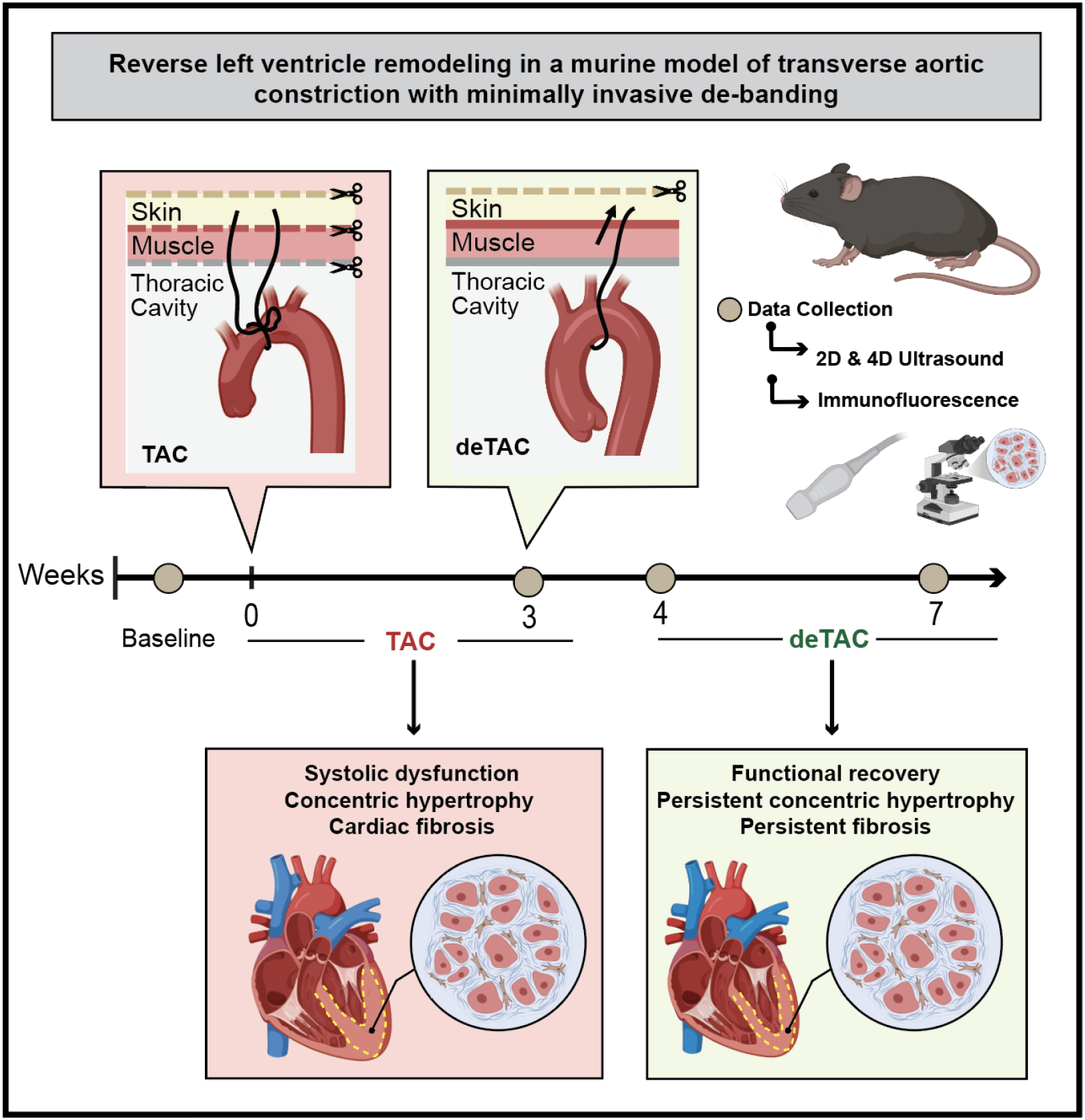

**HIGHLIGHTS:**

- Adaptation to the widely used transverse aortic constriction (TAC) procedure allows for minimally invasive de-TAC for studying reverse remodeling.
- This model increases feasibility and reproducibility, with no requirement for additional instruments or materials.
- State-of the art 4DUS assessment of cardiac function, mechanics, and geometry shows extensive left-ventricular remodeling and significant recovery of function following de-TAC.
- Cardiac fibrosis is persistent following functional recovery from heart failure with reduced ejection fraction (HFrEF).

## INTRODUCTION

Heart failure is the leading cause of death, affecting more than 23 million people worldwide.^1^ Pressure overload commonly occurs due to pathologic conditions such as hypertension or aortic stenosis^2^ and may lead to de-compensated heart failure when left untreated.^3,4^ Interestingly, the heart demonstrates an incredible potential for pathological resilience when treated properly for heart failure with reduced ejection fraction (HFrEF).^5^ At a molecular level, cardiac recovery from HFrEF in humans has recently been associated with decreased type I collagen synthesis, as demonstrated by decreased levels of procollagen type I C-terminal pro-peptide.^6^ Similarly in patients with left ventricular assist devices (LVAD), recovery from HFrEF has been linked to a dynamic response of interstitial cells, notably fibroblasts and macrophages.^7^ This functional and structural normalization occurs via a process known as reverse remodeling.^8^ However, the underlying mechanisms of functional and tissue-level recovery remain poorly understood, leaving blind spots in our understanding of the therapeutic mechanisms that promote recovery. Consequently, the development of animal models that accurately mimic human disease and reverse remodeling remains critical for elucidating the cellular and molecular mechanisms that drive functional recovery.

Transverse aortic constriction (TAC) in mice, first introduced by Rockman *et al*., is a common model of pressure overload that induces LV remodeling and functional decline. In the past decade, “deTAC” murine models of pressure overload that incorporate a recovery period have provided valuable insights into the mechanisms of cardiac dysfunction and recovery, and the potential for therapeutic interventions.^9–12^ However, most current deTAC models, which involve aortic banding followed by subsequent band release, are difficult to implement. Debanding in these models often requires a second a high-risk procedure such as thoracotomy or sternotomy,^13,14^ increasing the risk of bleeding, death, and post-surgical inflammation. These surgical risks may compromise the reliability of experimental outcomes. The use of absorbable sutures^15,16^ has previously been leveraged to circumvent the need for surgical band removal. However, such studies require more frequent imaging to control and detect the approximate point in time at which rapid aortic expansion or de-banding occurs. Furthermore, absorbable sutures are limited in their utility when a controlled experimental timeline is desired.

To address these challenges and limitations, we developed a novel and highly reproducible adaptation to the TAC procedure that enables deTAC/aortic de-banding using non-absorbable sutures and without the need for re-opening the thoracic cavity. Our refined method provides greater temporal control and ease of implementation, enabling more precise and enhanced studies of cardiac pressure overload and recovery dynamics. Using high-resolution two-dimensional and four-dimensional ultrasound, we characterized the functional and biomechanical recovery, and fibrotic response of the left ventricle (LV) following the resolution of pressure overload.

## METHODS

### Animal Information

All procedures were abided by the European Directive (2010/63/EU) and French laws for laboratory animal use and were approved by the local ethics committee (CCEA n°36; protocol no. 2023013113485359). We followed the ARRIVE guideline 2.0 for animal reporting.^17^ Aseptic technique was used for all surgical procedures.

### Surgical Method: Transverse aortic constriction with closed-chest de-banding

The TAC and deTAC procedures were performed by an experienced animal surgeon. We surgically induced transverse aortic constriction (TAC) with minimally invasive de-banding (deTAC) in 8-week-old C57Bl6/J (Janvier labs, France) male mice (n=10). We collected whole hearts from age-matched shams (n=4) at baseline, and from age-matched permanent TAC mice at 3W TAC (n=4) and 7W TAC (n=3).

#### General Anesthesia and Physiologic Monitoring

We anesthetized animals with 3% isoflurane in an induction chamber and then transferred them to a nose cone on a heating pad. Body temperature was monitored with a rectal thermometer and maintained at 37°C. ECG signals were monitored via electrodes throughout experimental procedures (PowerLab 26T, AdInstruments, U.K.).

#### Transverse aortic constriction

1- Animals were endotracheally intubated, removed from the nosecone, and placed on a ventilator (MiniVent Type 845, Hugo Sachs Elektronik, Germany) at 2% isoflurane with a tidal volume of 225 µL and a respiratory rate of 140 breaths/min. After performing a left-sided thoracotomy in the second intercostal space, the ribs were retracted to expose the transverse aorta. A 6-0 PVDF (Polyvinylidene fluoride) suture was placed underneath the aorta between the brachiocephalic and left common carotid artery. Representative images of the surgical procedure are available in **Supplemental Figure S1**.
2- A 27-G blunt-tip needle was positioned parallel to the anterior surface of the transverse aorta. The 6-0 PVDF suture was tied around the blunted needle using a mooring hitch knot (**Fig. S2)**, ensuring a secure loop around the needle. Immediately after creating the knot, the blunt needle was removed. With the ends of the constriction suture extending through the muscle layer (**Fig. 1A**), the thoracic cavity and skin were closed using a 5-0 resorbable suture. After completing the procedure, in vivo high-frequency ultrasound was performed to confirm the placement of the constriction.
3- Animals were extubated and placed in a clean cage on a warm pad. We subcutaneously administered buprenorphine (1mg/kg) twice per day for pain management.

**Figure 1.**
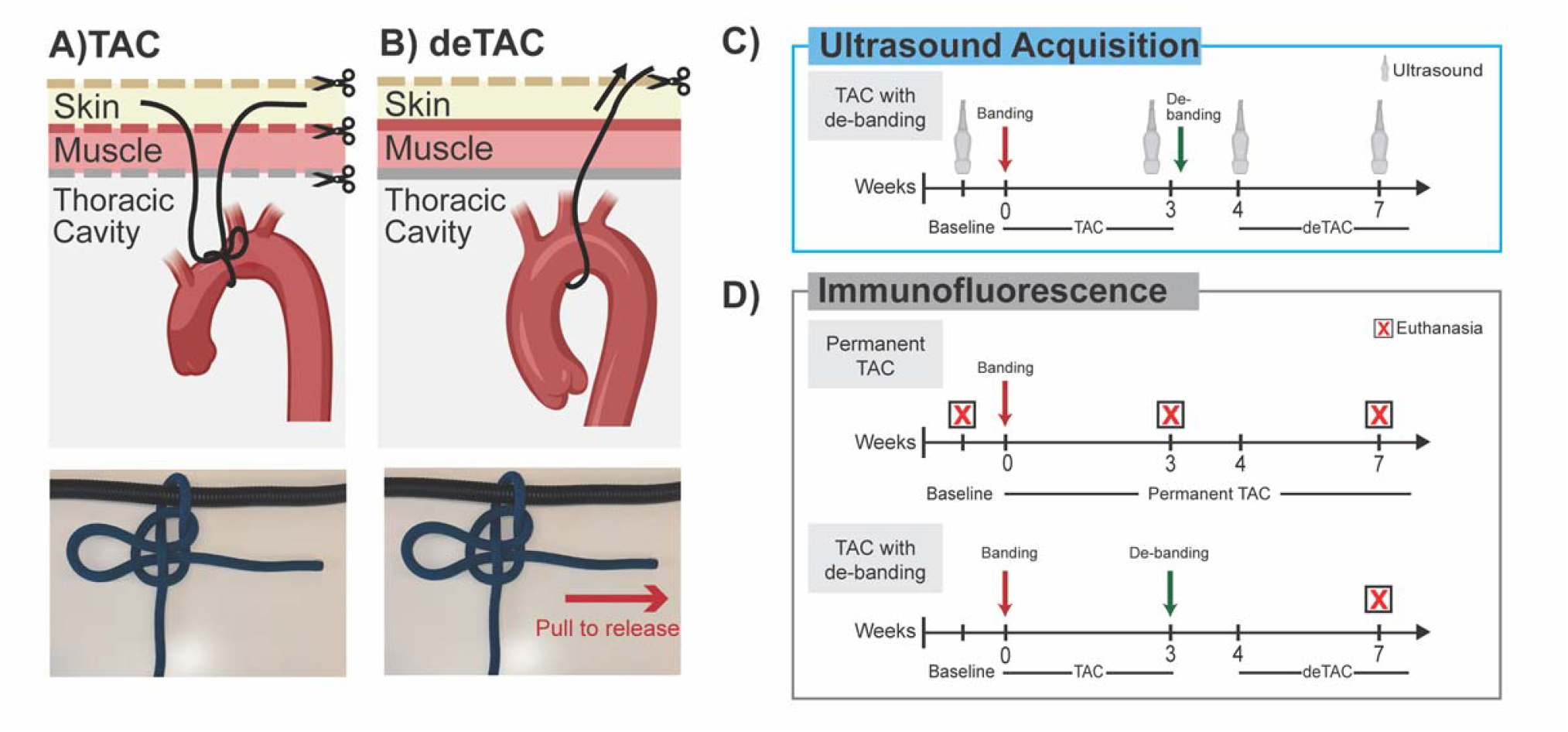
Experimental Overview: Schematic of **A)** TAC and **B)** minimally invasive deTAC. Black arrows = direction in which suture ends are pulled. Scissor icon=incised tissue layer. Timelines for **C)** ultrasound acquisition and **D)** immunofluorescence. W= week, TAC=transverse aortic constriction, deTAC= aortic de-banding. Created in Biorender.

#### Closed chest aortic de-banding (deTAC)

4- Aortic de-banding (deTAC) was performed at three weeks after TAC or after EF was less than 45% when measured by 2D ultrasound (**Fig. 1B**). Animals were anesthetized and the skin layer above the left-sided third intercostal space was reopened to expose the two ends of the 6-0 PVDF suture. By gently pulling one end of the suture with forceps, the mooring hitch knot was released from the aorta, allowing for the closed-chest de-banding procedure (**Fig. S1-2**). The skin was then closed with a 5-0 resorbable suture. From start to finish, the de-banding procedure takes 5 minutes.
5- Immediately after completing the procedure, in vivo high-frequency ultrasound was performed to confirm the de-banding of the aorta.

### In Vivo High-Frequency Ultrasound

All ultrasound images were acquired with the Vevo 3100 ultrasound system (FUJIFILM VisualSonics) using the MX550D linear array transducer (40 MHz center frequency). We collected ultrasound images of the transverse aorta and LV at baseline prior to inducing TAC, 3 weeks (W) after TAC, 1W after deTAC, and 4W after deTAC (**Fig. 1C**).

After anesthetizing the animals, we used depilatory cream to remove the fur from the chest wall. We collected B-mode images of the transverse aorta and pulsed-wave Doppler (PWD) downstream from the constriction site to verify the surgical procedure. Elevated aortic valve velocity and visual narrowing of the transverse aorta were used to confirm successful banding (TAC) and the reverse for de-banding (deTAC). To assess cardiac function and wall geometry, we collected long-axis (LAX) and short-axis (SAX) B-mode images, and SAX M-mode images of the LV. We acquired a 4D (3D+time) image of the LV to quantify mass and kinematics. We collected 4D images using a linear stop motor (step size= 0.13 mm) to scan across the SAX from the base of the apical epicardium to the ascending aorta.

### Image Analysis

#### 2D Image Analysis

All 2D image analysis was performed in VevoLab (Version 3.9.0, FUJIFILM VisualSonics). We measured aortic valve velocity using peak velocity traces of PWD images and ejection fraction (EF), stroke volume (SV), and cardiac output (CO) from 2D B-mode images. LV wall thickness, measured as the average of posterior and anterior wall thickness, and diameter were measured from SAX M-mode images at the center of the ventricle during diastole.

#### Four-Dimensional Analysis

We performed 4D strain analysis of the LV at each timepoint using a custom-built MATLAB graphical user interface as previously described.^18,19^ In brief, 4D images were aligned with the centerline of the LV in the long axis and segmented throughout the cardiac cycle. Segmentation points were placed at four equally spaced short-axis segments from base to apex, with six equally spaced points around the endo- and epi-cardial wall of each slice. The remaining myocardial points were then spatially and temporally interpolated from the segmented points and used to create a dynamic 3D mesh from which LV volume and mass, and circumferential-, longitudinal-, radial-, and surface area strain were calculated. All strains were calculated using a Lagrangian reference frame and the equations for small engineering strain as previously described.^19^ To obtain localized strain estimates, we generated strain curves at evenly divided slices across the ventricle. Global peak strains were calculated by averaging the strains across each respective region. Heat maps color schemes were generated using the Light Bartlein Color Maps^20^ add-on in MATLAB.

### Histology & Immunohistochemistry

Whole hearts were collected from sham (n=4), permanent-TAC animals at 3W TAC (n=4) and 7W TAC (n=3), and from deTAC animals at 4-week deTAC (n=4) (**Fig. 1D**). Hearts were fixed overnight in 4% PFA, subjected to a sucrose gradient (5%, 10%, 20% sucrose in PBS) and included in OCT. 12µm sections were washed (PBS), permeabilized (PBS, 0.1% Triton), blocked in blocking solution (BS) (PBS, 0.1% Triton, 2% BSA) and incubated with primary antibody (Goat anti-Collagen type I, Southern Biotech 1310-01; Goat anti-PDGFRα, R&D, AF1062) overnight. Sections were then washed (PBS) and incubated for 2 hours with secondary antibody in BS (Anti-Goat, Alexa 647, ThermoFisher A32849) and DAPI. Sections were then washed (PBS) and mounted (Fluoromount, Sigma). Images were acquired with a ZEISS Axioscan 7 and were processed with Zen and ImageJ software.^21^ Collagen Type I^+^ area was measured in two to three consecutive sections per individual heart. PDGFRα^+^ fibroblasts were counted in five fields in the LV and septum from equivalent sections from each animal.

### Statistical Analysis

We performed statistical analysis in Prism 10 (GraphPad Software, San Diego, California, USA) with *p<0*.*05* indicating significance. All data were tested for normal distribution using the Shapiro-Wilk test and were reported as mean ± standard deviation. For normally distributed data we performed a repeated measure analysis of variance (ANOVA) with post hoc Tukey’s multiple comparisons test to compare cardiac function, geometry, and global strain values across baseline, 3W TAC, 1W deTAC, and 4W deTAC. We used a two-way repeated measures ANOVA with a post-hoc Tukey’s multiple comparisons test to assess regional differences in strain. A Friedman test with Dunn’s multiple comparisons test was performed for data that were not normally distributed. For histological examinations, we used one-way ANOVA with Tukey’s post-hoc test.

## RESULTS

### TAC and closed-chest deTAC Surgical Procedure

The refined murine model of the reversible aortic constriction method is detailed in the Methods section. A schematic experimental overview (**Fig.1**) is provided to illustrate the key steps of the closed-chest de-banding procedure (**Fig. S1 and S2**). During this procedure, two out of ten animals died.

### Verification of TAC and deTAC

We observed both anatomical and hemodynamic indications after TAC. The TAC procedure induced visible narrowing of the transverse aorta that was detectable in the sagittal view with ultrasound (**Fig. 2A**). We verified the severity of the TAC procedure with pulsed-wave Doppler and observed a significant elevation of the flow velocity peak at the constriction site after surgery. Blood flow velocity remained elevated until the 3W TAC timepoint (Baseline: 980 ± 121 mm/s, 3W TAC: 3738 ± 396 mm/s, *p*<0.001) (**2B-C, Table S1**). After three weeks of pressure overload our refined suture technique allowed us to remove the aortic constriction without a second thoracotomy. This resulted in a decrease of flow velocity peak at 1W- (1781 ± 444 mm/s, *p*=0.317) and 4W- (1823 ± 712 mm/s, *p*=0.040) deTAC compared to 3W TAC (**Fig 2B-C**).

**Figure 2.**
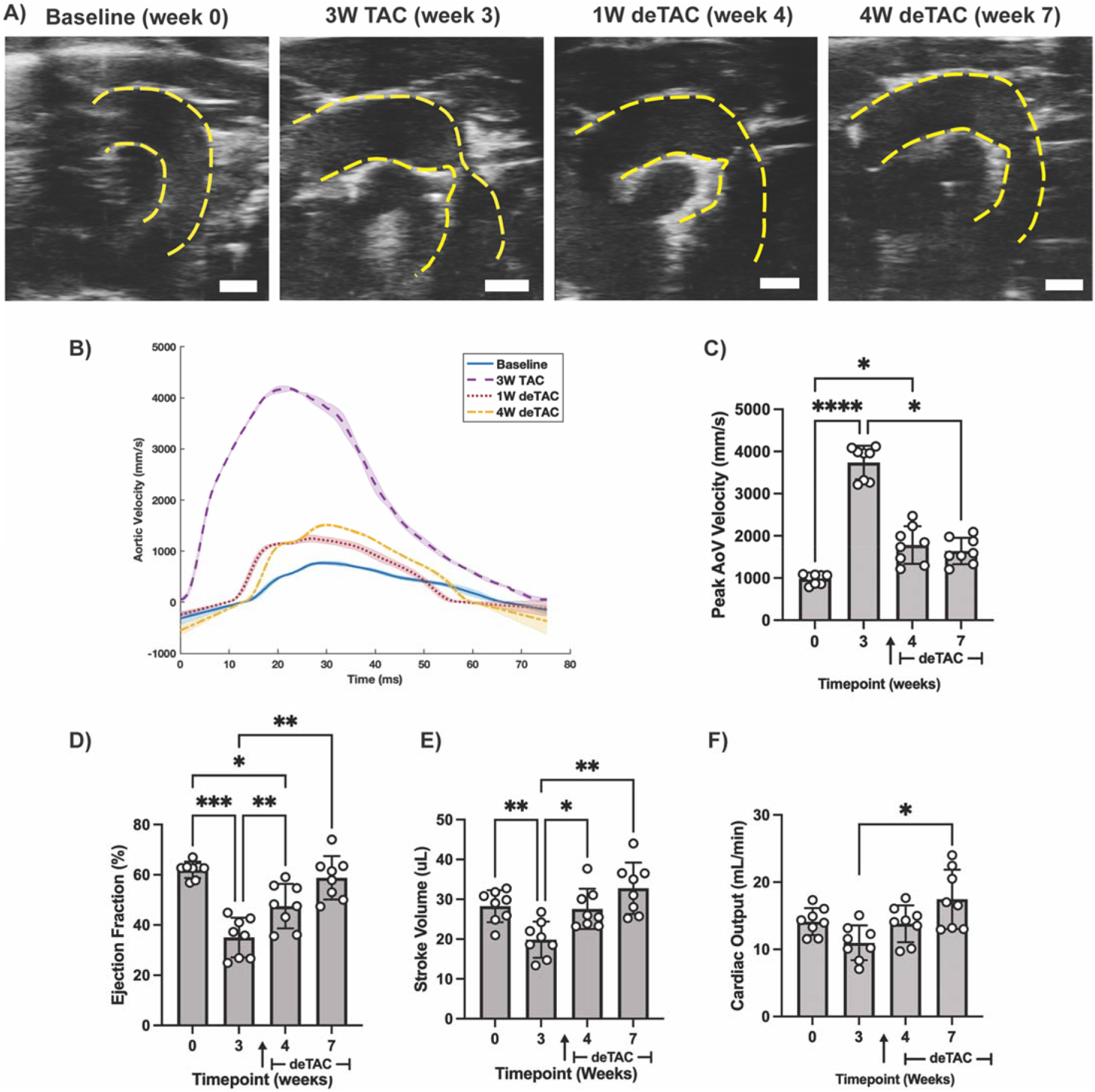
Verification of Procedure and Cardiac Function: **A)** Sagittal B-mode ultrasound images of the transverse aorta (outlined in yellow). **B)** Representative aortic velocity from a single animal (n=1), where the shaded region is the standard deviation of 3-5 waveforms, and **C)** peak aortic velocity measured from pulsed-wave Doppler. **D)** Ejection fraction, **E)** stroke volume, and **F)** cardiac output measured from PSLAX ultrasound. Scale bar= 1mm. \**p<*0.05, \*\**p*<0.01, \*\*\**p*<0.001, \*\*\*\**p*<0.0001. Normally distributed data was analyzed using repeated measures ANOVA with Tukey’s post hoc test. Non-parametric equivalents were used for non-normally distributed data. n=8/group.

### Refined TAC-deTAC model induce heart failure with functional recovery

After TAC, animals developed significant systolic dysfunction (EF<45%). Both EF and SV significantly decreased at 3W TAC compared to baseline (EF: 35.0 ± 7.9%, *p*<0.001; SV: 19.9 ± 4.5 uL, *p*=0.003) (**Fig. 2D-E**). Following de-banding, EF and SV showed progressive sustained recovery throughout the deTAC period. EF increased to 47.5 ± 8.8% (*p*= 0.003) by 1W deTAC and to 58.8 ± 8.6% by 4W deTAC (*p*=0.002). SV increased to 27.6 ± 5.1 uL (*p*=0.020) at 1W deTAC and 32.8 ± 6.5 uL at 4W deTAC (*p*=0.006) (**Fig. 2E**). CO decreased during TAC compared to baseline, though not significantly (Baseline: 14.1 ± 2.0 mL/min, 3W TAC: 11.0 ± 2.6 mL/min, *p*=0.052) and increased after de-banding at 1W- (13.8 ± 2.7 mL/min, *p*=0.084) and 4W-deTAC (17.5 ± 4.4 mL/min, *p*=0.022) compared to 3W TAC (**Fig. 2F**). Values are summarized in **Supplemental Table 1**.

Using 4D strain analysis, we assessed whether TAC and reverse TAC led to any LV regional or global strain modifications. During TAC, global and regional LV strain metrics showed pronounced alterations that aligned with adverse remodeling observed in 2D ultrasound (**Fig. 4; Fig S3-6**). Peak global circumferential (E_cc_), longitudinal strain (E_ll_), radial (E_rr_), and surface area strain (E_sa_) all declined substantially at 3W TAC (E_cc_: -17.7 + 4.0%, p=0.002; E_ll_: - 11.4 ± 3.3%, p=0.006; E_rr_: 12.6 ± 4.5%, p=0.018; E_sa_= -28.8 ± 6.8%, *p*<0.001) (**Fig. 3**). We observed minor heterogeneity in radial- and surface area strain after 3W TAC, as indicated by interlaced areas of orange and blue in heatmaps (**Fig S3-S6**). After deTAC, global LV strain showed rapid recovery, as indicated by the return of orange in the heatmaps (**Fig. 3**). These parameters exhibited progressive improvement at 1W deTAC (E_cc_: -23.4 ± 5.2%, p=0.002 ; E_ll_: - 16.8 ± 3.4%, p=0.052; E_rr_: 18.0 ± 3.2%, *p*=0.029; E_sa_: -39.6 ± 8.2%, *p*=0.005) and 4W deTAC (E_cc_: -17.1 ± 2.0%, *p*=0.001 ; E_ll_: -17.1 ± 2.0%, p=0.001; E_rr_: 16.4 ± 5.2%, p=X; E_sa_: -41.8 ± 5.5%, *p*=0.526). Interestingly, heat maps showed that reverse TAC induced segmental recovery in radial- and surface area strain. However, we did not identify a distinct and specific regional pattern dependent on LV reversibility (**Fig S3-6**). All global strain values are reported in **Table S1**. Altogether, these findings suggest that our refined TAC-deTAC model induces heart failure with functional recovery.

**Figure 3.**
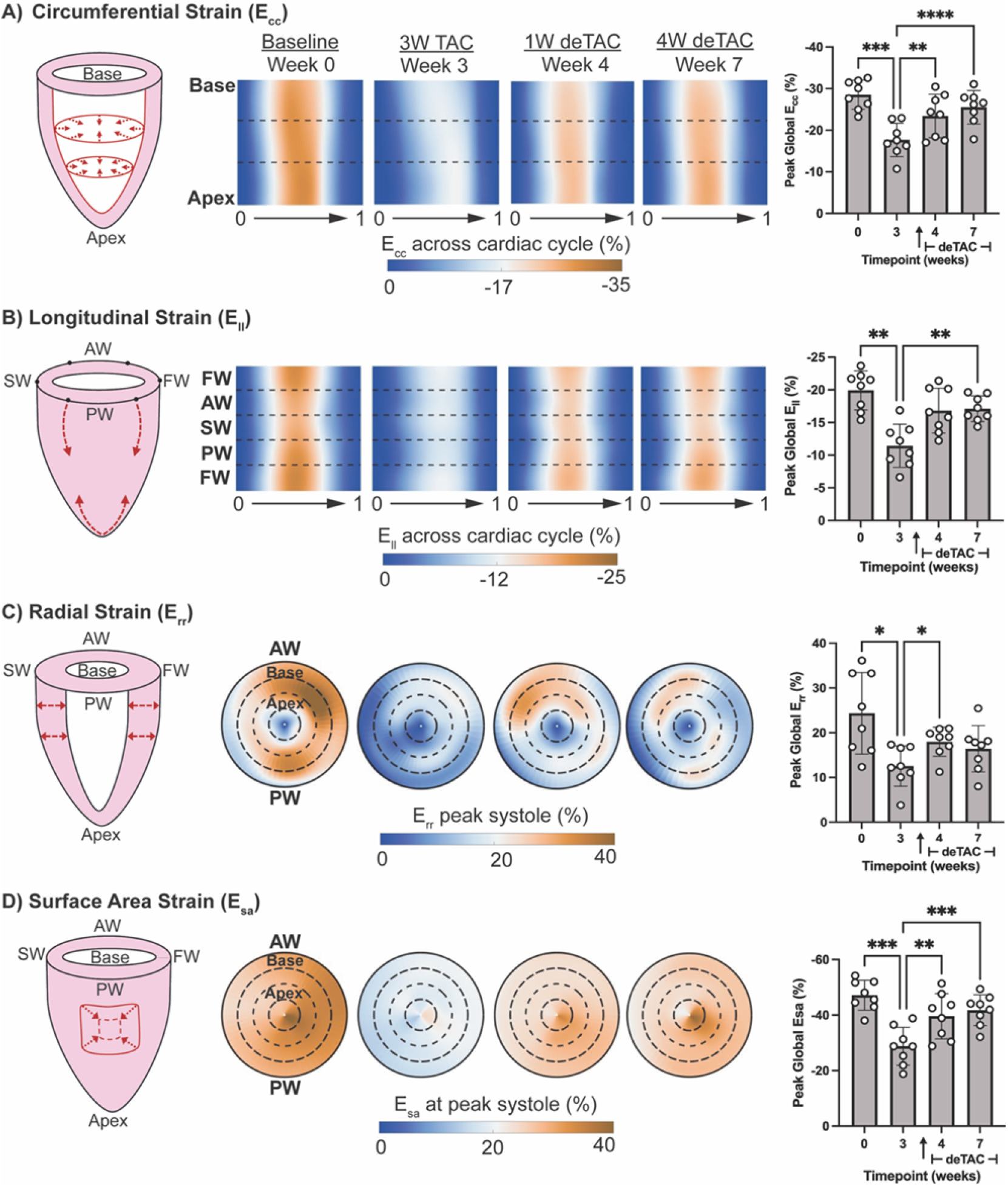
Left Ventricle Strain. Schematic of strain annotated with red dashed arrows, average heatmaps, and peak global strains for **A)** Circumferential strain (E_cc_) **B)** Longitudinal Strain (E_ll_), **C)** Radial Strain (E_rr_), **D)** Surface area strain (E_sa_). FW= free wall, AW=anterior wall, SW=septal wall, PW=posterior free wall. \**p<*0.05, \*\**p<*0.01, \*\*\**p<*0.001. Data was analyzed using repeated measures ANOVA with Tukey’s post hoc test. n=8/group.

**Figure 4.**
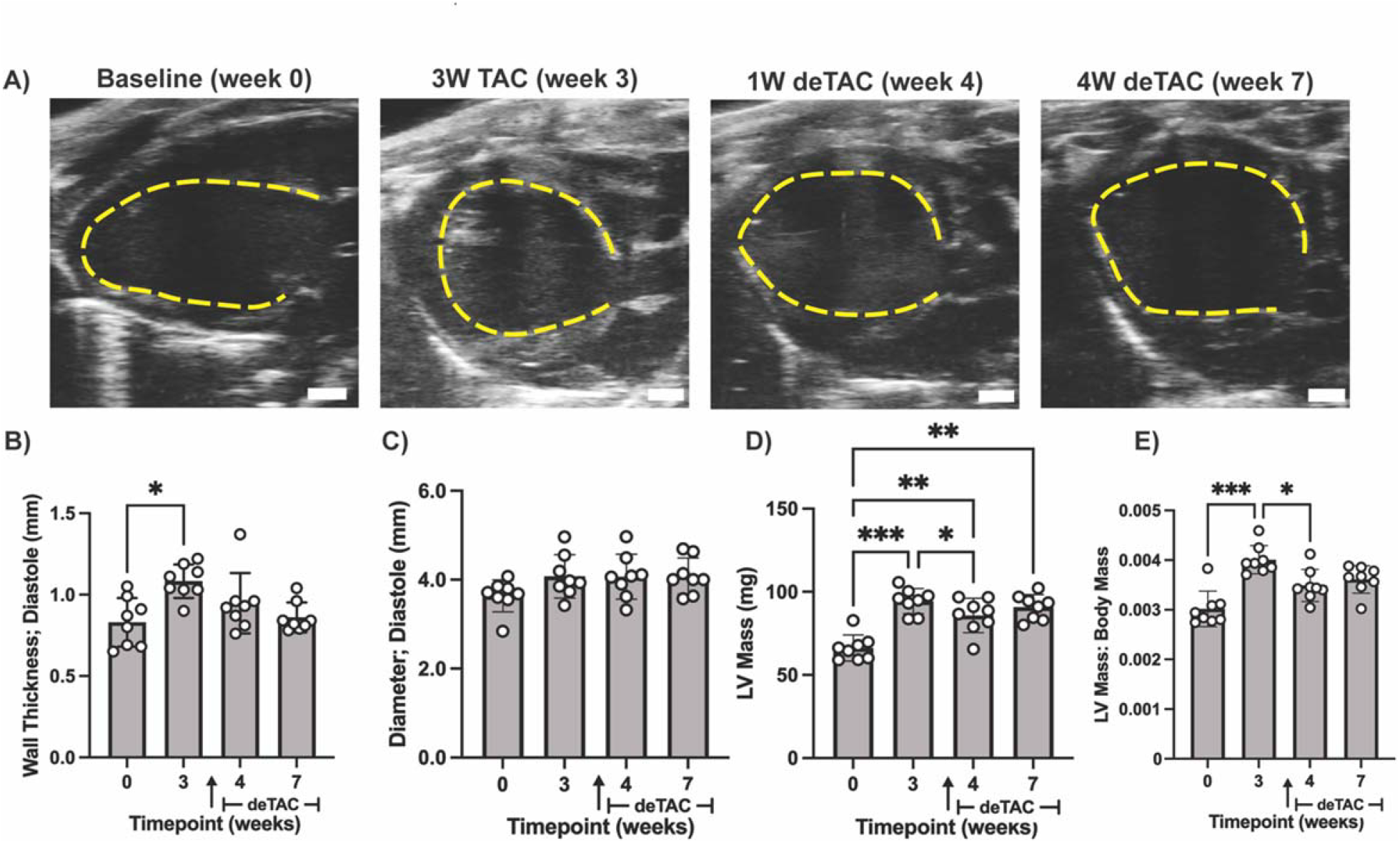
Left ventricle geometry. Representative **A)** Parasternal long axis B-modes of the left ventricle (LV) with endocardium demarcated with dotted line. Scale bar= 1mm. **B-C)** M-mode measured diastolic wall thickness and diameter, respectively. **D-E)** 4D ultrasound measurements of left ventricle (LV) mass. Dia.=diastole. \**p<*0.05, \*\**p<*0.01, \*\*\**p<*0.001. Normally distributed data was analyzed using repeated measures ANOVA with Tukey’s post hoc test. Non-parametric equivalents were used for non-normally distributed data. n=8/group.

### deTAC did not result in myocardial structural remodeling despite functional recovery

To further characterize myocardial structural remodeling in our TAC-deTAC procedure, we used a combination of 2D/4D ultrasound and histological techniques to assess the trajectory of LV hypertrophy and fibrosis. We observed geometric remodeling in the LV during the TAC period, with partial reverse remodeling after deTAC (**Table S1**). Wall thickening and increased sphericity was visible in PSLAX B-mode at 3W TAC relative to baseline (**Fig 4A**). This change in ventricle shape was accompanied by a significant increase in wall thickness (Baseline: 0.8 ± 0.2 mm, 3W TAC: 1.1 ± 0.1 mm, *p*=0.011). Wall thickness slightly recovered towards baseline after aortic de-banding, though the decrease was not significant (1W deTAC: 0.9 ± 0.2mm, *p*=0.728; 4W deTAC: 0.9 ± 0.1mm, *p=*0.120) (**Fig 4B**). In contrast to wall thickness, LV diastolic diameter remained relatively unchanged between experimental timepoints, ranging from 3.6 ± 0.4 mm to 4.1 ± 0.4 mm (**Fig. 4C**). LV mass and relative LV mass (LV mass/body mass) both significantly increased after 3W TAC. LV mass remained elevated, while relative LV mass only partially decreased at 1W deTAC and a plateaued at 4W deTAC (**Fig. 4D-E**).

Lastly, we quantified Collagen Type I deposition in the LV (**Fig. 5A-C**). We observed a significant increase in the percentage of collagen type I^+^ area at 3W TAC (7.7 ± 2.8%, n=4, *p*=0.029.) and 7W TAC (8.2 ± 4.2%, n=4, *p*=0.018) compared to baseline (1.4 ± 0.5%, n=4) (**Fig. 5C**). At 4W deTAC, despite functional recovery and structural normalization, Collagen type I^+^ area remained elevated (7.2 ± 2.1%, n=4, *p*=0.034) compared to baseline, and did not significantly differ from TAC. Hence, cardiac fibrosis did not drop significantly in deTAC animals.

**Figure 5.**
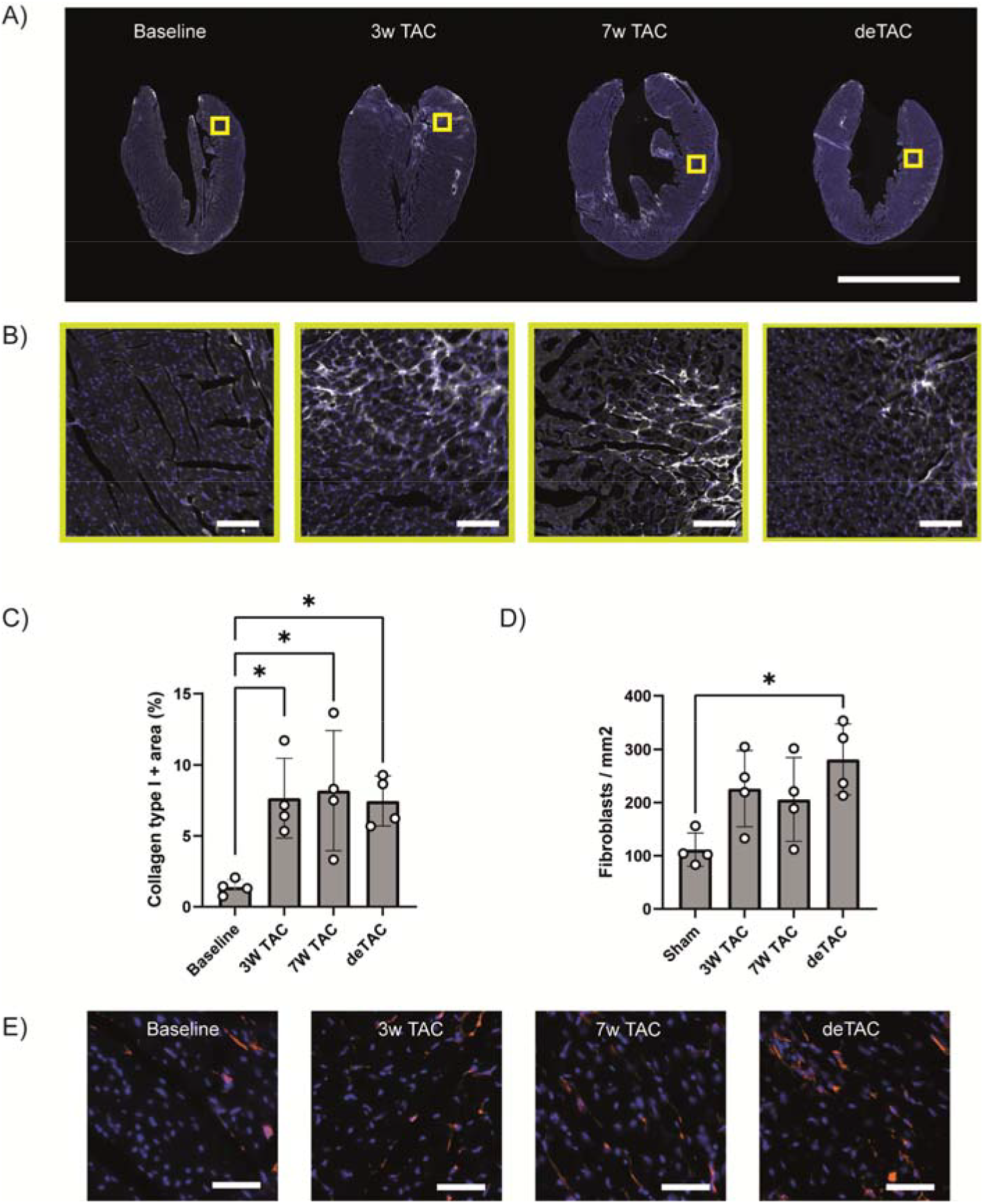
ECM content and fibroblast numbers remain elevated during functional recovery. **A)** Representative 4-chamber view of control, 3 weeks TAC, 7 weeks TAC, and 3 weeks TAC + 4 weeks deTAC. Immunofluorescence staining shows collagen type I (grayscale) and DAPI (blue). Scale bar is 5mm. **B)** Insets form the images in A) Scale bars are 100µm. **C)** Quantification of Collagen type I+ area over total area in the left ventricle and septum shows a significant increase in TAC and RR compared to baseline. **D)** Quantification of the numbers of PDGFRα+ fibroblasts in the LV/septum. **E)** Representative immunofluorescence images showing PDGFRα+ fibroblasts. Scale bar 50µm. \**p<0*.*05*, ANOVA with Tukey’s post hoc test, n=4/group.

Cardiac fibroblasts are the main cell-type responsible for Collagen type I production and turnover. Hence, we quantified the numbers of PDGFRα^+^ cardiac fibroblasts in sections adjacent to those used for measuring Collagen type I^+^ area (**Fig. 5D, E**). Interestingly, compared to baseline (111.4 ± 31.3 fibroblasts/mm^2^, n=4), we found the most elevated numbers of fibroblasts in the deTAC hearts (281.1 ± 67 fibroblasts/mm^2^, n=4, *p*=0.014), with lower numbers of fibroblasts at 3W (226.2 ± 71.7 fibroblasts/mm^2^, *p*=0.110) and 7W TAC (205.9 ± 78.8 fibroblasts/mm^2^, n=4, *p*=0.220) (**Fig. 5D**). Hence, in our model, significant functional recovery was not associated with reduced fibrosis or cardiac fibroblast numbers.

## DISCUSSION

In this study, we demonstrate the feasibility of a refined aortic de-banding method that enables the investigation of recovery from heart failure without the need to reopen the chest cavity. In our longitudinal ultrasound study, we confirmed that this new minimally invasive method induces sustained systolic dysfunction (EF<45%) with cardiac reverse remodeling after de-banding, hence reproducing reverse remodeling similar to that observed in heart failure patients with recovered EF. By limiting open surgical intervention, our approach represents an accessible and advantageous adaptation to an already well-established TAC model. This will be particularly useful for studying the effects of mechanical unloading at early timepoints following the afterload increase from TAC surgery.^9,11^ While our study specifically induced HFrEF,^22,23^ this model could easily be modified to study RR after HFpEF^24^ by adjusting the degree of initial aortic constriction.^25,26^ Additionally, the use of non-resorbable sutures allows for precise control over the de-banding timepoint. Thus, sustained pressure overload can occur over a designated period and can be confirmed via measurement of sustained elevated blood flow velocity. This increases reproducibility, allowing for reduced number of animals to be used.

As expected, we observed a decrease in cardiac function with pressure overload similar to permanent TAC models.^26–29^ EF decreased below 45% following banding, indicating systolic dysfunction, and gradually normalized to >55% after de-banding suggesting that our model is suitable for studying recovery from heart failure with reduced ejection fraction (HFrEF). The dramatic decrease in strain, further confirmed the presence of systolic dysfunction following the TAC procedure. Additionally, the unchanged LV cavity diameter coupled with increased myocardial wall thickness suggests that the heart experienced concentric hypertrophy during the TAC period. Despite substantial decreases in cardiac function (EF, SV, CO) and strain, the LV demonstrated significant recovery towards baseline after de-banding. Cardiac perfusion and global strain trended towards baseline in as little as one week after deTAC. Cardiac hypertrophy appeared nonreversible as LV wall thickness, diameter, and mass did not return to baseline following deTAC, potentially due to persistent fibrosis. Reversal of concentric hypertrophy has been seen in some deTAC models following less severe pressure overload,^13^ suggesting that the heart’s ability to geometrically recover may depend on the severity of pressure overload and highlighting the importance of early diagnosis to increase the probability of cardiac recovery.

Fibrosis is a major driver of heart failure progression.^30^ The relationship between RR and fibrosis remains controversial, and likely depends on the type and degree of heart failure, and time allowed for RR to occur.^31^ Recovery from HFpEF in mice, characterized by normalization of diastolic dysfunction, has been associated with a regression of fibrosis.^32^ Additionally, aortic de-banding in mice with HFrEF has been seen to result in reduced transcription of *Col1a1*, which encodes for type I collagen.^10^ We found that fibrosis persisted up to one month after removal of the aortic constriction. This is in line with clinical studies with histological analysis revealing elevated fibrosis despite reduced collagen expression following mechanical unloading in HFrEF patients.^33^ In order to further characterize fibrosis, we quantified the relative numbers of cardiac fibroblasts, the main cell-type responsible for extracellular matrix (ECM) deposition. Following deTAC, cardiac fibroblast numbers remained elevated, further supporting the presence of fibrosis despite mechanical unloading. Single-nucleus RNAseq-based analysis of LVAD patient biopsies has revealed that fibroblasts have a dynamic transcriptional response to mechanical unloading.^7^ Characterization of ECM quality (e.g. level of crosslinking) and the nature of cardiac fibroblast activity in RR will help to clarify the complex relationship between myocardial fibrosis and functional recovery in our model.

Our refined TAC-deTAC method incorporates straightforward yet advantageous adaptations to the well-established murine TAC model. Notably, the degree of invasive intervention required in our model and the immediate significant opening of the aorta more closely mimic those of current minimally-invasive HF treatments such as percutaneous valve implantation^34^ and minimizes confounding factors. This model will be a valuable tool for studying the functional and molecular changes that occur during left ventricular loading and unloading. It holds promise for advancing research on the mechanisms driving reverse remodeling, which could reveal potential therapeutic targets for heart failure.

## Study Limitations

While our refined TAC-deTAC model provides a valuable method for studying reverse recovery from pressure overload, several limitations should be considered. First, fibrotic tissue around the suture resulting from prolonged constriction can make suture removal challenging. After three weeks of TAC, the success rate for suture removal decreases significantly to less than 50% and continues to decline with longer periods of TAC. To address this issue in future studies, sutures of varying gauge, material, or coating may be considered. Second, we did not compare our approach to classical deTAC methods to evaluate potential differences in inflammation or other postoperative outcomes. However, our refine deTAC method only requires five minutes to be performed and assumes reduced risk that aims to minimize the number of animals used for additional survival studies. Other studies, such as myocardial infarction models involving second survival procedures, provide relevant comparisons.^35^ Lastly, our study utilized the well-established model of TAC induces an immediate increase in afterload that is not reflective of the gradual progression of aortic constriction in patients with longstanding cardiovascular disease.^36^ The rapid onset of pressure overload of our model may influence short-term hemodynamic measurements and warrants further investigation. We would encourage future studies examining different rates of insult to implement modified versions of our deTAC.

## CONCLUSION

In conclusion, we have designed and validated a refined methodology for studying heart failure reverse remodeling. This approach utilizes a modified, releasable knot for aortic constriction in a murine model of pressure overload. Our method provides a valuable tool for both fundamental and translational research, effectively mimicking conditions such as aortic stenosis and therapeutic interventions like aortic valve implantation.

## CLINICAL PERSPECTIVES

### Competency in Medical Knowledge

Heart failure with reduced ejection fraction (HFrEF) continues to pose significant clinical challenges with poor prognosis and limited treatment options. Promisingly, functional and structural recovery has been observed in a subset of aortic stenosis patients after transverse aortic valve replacement (TAVR) and surgical aortic valve replacement (TAVI).^37–40^ However, the underlying mechanisms of this reverse remodeling remain incompletely understood, making it difficult to develop treatments that target recovery. This is largely due to the lack of accurate and reproducible experimental models. By being able to control the timing of release, minimizing surgical invasion, and allowing for immediate relief of pressure overload, our method introduces a clinically relevant model of recovery from valve replacement or aortic stenosis. Our model may be used to study and identify insights into the cellular and molecular mechanisms driving functional and structural recovery after treatment of pressure overload.

### Translational Outlook

The development of therapeutic strategies to promote recovery from heart failure requires experimental models that accurately reflect the dynamic nature of cardiac remodeling. Our refined TAC-deTAC model offers a practical and reproducible approach for studying RR, eliminating the need for a second invasive surgery while maintaining precise control over pressure overload release. This advances the ability to investigate the timing and mechanisms of RR, which would in turn allow us to design new therapeutic targets that promote cardiac function. Furthermore, our method more faithfully mimics HF treatment methods such as valve implantation surgery, which do not require opening the chest.^34^ Given that fibrosis persists despite improvements in cardiac perfusion, strain, and function, our model provides a platform for testing novel anti-fibrotic and regenerative therapies. By bridging the gap between preclinical research and clinical applications, this model has the potential to accelerate the development of targeted interventions aimed at improving long-term outcomes for patients with HFrEF.

## Acknowledgements

We gratefully thank the staff for animal housing (PhyMedExp, RAM) and would like to acknowledge Salomé Bélec from Imagerie du Petit Animal de Montpellier (IPAM) for her technical support and assistance with high-resolution ultrasound (LRQA Iso9001; France Life Imaging (grant ANR-11-INBS-0006); IBISA; Leducq Foundation (RETP), I-Site Muse).

## REFERENCES

1. Bui AL, Horwich TB, Fonarow GC. Epidemiology and risk profile of heart failure. Nat Rev Cardiol. 2011;8(1):30–41. doi:10.1038/nrcardio.2010.165

2. Shimizu I, Minamino T. Physiological and pathological cardiac hypertrophy. J Mol Cell Cardiol. 2016;97:245–262. doi:10.1016/j.yjmcc.2016.06.001

3. Mills KT, Bundy JD, Kelly TN, et al. Global Disparities of Hypertension Prevalence and Control: A Systematic Analysis of Population-based Studies from 90 Countries. Circulation. 2016;134(6):441–450. doi:10.1161/CIRCULATIONAHA.115.018912

4. Calderone A. Myocardial Hypertrophy and Regeneration. In: McManus LM, Mitchell RN, eds. Pathobiology of Human Disease. Academic Press; 2014:580–592. doi:10.1016/B978-0-12-386456-7.02107-9

5. Rodrigues PG, Leite-Moreira AF, Falcão-Pires I. Myocardial reverse remodeling: how far can we rewind? Am J Physiol-Heart Circ Physiol. 2016;310(11):H1402–H1422. doi:10.1152/ajpheart.00696.2015

6. Ravassa S, Lupón J, López B, et al. Prediction of Left Ventricular Reverse Remodeling and Outcomes by Circulating Collagen-Derived Peptides. JACC Heart Fail. 2023;11(1):58–72. doi:10.1016/j.jchf.2022.09.008

7. Amrute JM, Lai L, Ma P, et al. Defining cardiac functional recovery in end-stage heart failure at single-cell resolution. Nat Cardiovasc Res. 2023;2(4):399–416. doi:10.1038/s44161-023-00260-8

8. Falcão-Pires I, Ferreira AF, Trindade F, et al. Mechanisms of myocardial reverse remodelling and its clinical significance: A scientific statement of the ESC Working Group on Myocardial Function. Eur J Heart Fail. 2024;26(7):1454–1479. doi:10.1002/ejhf.3264

9. Abe K, Sasano T, Soejima Y, Fukayama H, Maeda S, Furukawa T. Hypermethylation of Hif3a and Ifltd1 is associated with atrial remodeling in pressure-overload murine model. Sci Rep. 2025;15(1):2699. doi:10.1038/s41598-025-85382-8

10. Perera-Gonzalez M, Kiss A, Kaiser P, et al. The Role of Tenascin C in Cardiac Reverse Remodeling Following Banding–Debanding of the Ascending Aorta. Int J Mol Sci. 2021;22(4):2023. doi:10.3390/ijms22042023

11. Topkara VK, Chambers KT, Yang KC, et al. Functional significance of the discordance between transcriptional profile and left ventricular structure/function during reverse remodeling. JCI Insight. 2016;1(4). doi:10.1172/jci.insight.86038

12. Zhang X, Javan H, Li L, et al. A modified murine model for the study of reverse cardiac remodelling. Exp Clin Cardiol. 2013;18(2):e115–e117.

13. Bjørnstad JL, Skrbic B, Sjaastad I, Bjørnstad S, Christensen G, Tønnessen T. A mouse model of reverse cardiac remodelling following banding-debanding of the ascending aorta. Acta Physiol Oxf Engl. 2012;205(1):92–102. doi:10.1111/j.1748-1716.2011.02369.x

14. Merino D, Gil A, Gómez J, et al. Experimental modelling of cardiac pressure overload hypertrophy: Modified technique for precise, reproducible, safe and easy aortic arch banding-debanding in mice. Sci Rep. 2018;8(1):1–10. doi:10.1038/s41598-018-21548-x

15. Pölzl L, Hirsch J, Eder J, et al. Innate Reverse Remodeling Reveals Novel Treatment Option for Heart Failure. In: The Thoracic and Cardiovascular Surgeon. Vol 71. Georg Thieme Verlag KG; 2023:DGTHG-KV20. doi:10.1055/s-0043-1761741

16. Lao Y, Zheng C, Zhu H, Lin H, Huang X, Liao Y. Operating Transverse Aortic Constriction with Absorbable Suture to Obtain Transient Myocardial Hypertrophy. J Vis Exp JoVE. 2020;(163):e61686. doi:10.3791/61686

17. Percie du Sert N, Hurst V, Ahluwalia A, et al. The ARRIVE guidelines 2.0: Updated guidelines for reporting animal research. BMC Vet Res. 2020;16:242. doi:10.1186/s12917-020-02451-y

18. Damen FW, Berman AG, Soepriatna AH, et al. High-Frequency 4-Dimensional Ultrasound (4DUS): A Reliable Method for Assessing Murine Cardiac Function. Tomogr Ann Arbor Mich. 2017;3(4):180–187. doi:10.18383/j.tom.2017.00016

19. Earl CC, Pyle VI, Clark SQ, et al. Localized strain characterization of cardiomyopathy in Duchenne muscular dystrophy using novel 4D kinematic analysis of cine cardiovascular magnetic resonance. J Cardiovasc Magn Reson. 2023;25:14. doi:10.1186/s12968-023-00922-3

20. Light Bartlein Color Maps. January 16, 2025. Accessed January 16, 2025. https://www.mathworks.com/matlabcentral/fileexchange/17555-light-bartlein-color-maps

21. Schneider CA, Rasband WS, Eliceiri KW. NIH Image to ImageJ: 25 years of image analysis. Nat Methods. 2012;9(7):671–675. doi:10.1038/nmeth.2089

22. Weinheimer CJ, Lai L, Kelly DP, Kovacs A. A Novel Mouse Model of Left Ventricular Pressure Overload and Infarction Causing Predictable Ventricular Remodelling and Progression to Heart Failure. Clin Exp Pharmacol Physiol. 2015;42(1):33–40. doi:10.1111/1440-1681.12318

23. Sayour NV, Gergely TG, Váradi B, et al. Comparison of mouse models of heart failure with reduced ejection fraction. ESC Heart Fail. 2025;12(1):87–100. doi:10.1002/ehf2.15031

24. Valero-Muñoz M, Backman W, Sam F. Murine Models of Heart Failure With Preserved Ejection Fraction. JACC Basic Transl Sci. 2017;2(6):770–789. doi:10.1016/j.jacbts.2017.07.013

25. Deng J, Li D, Zhang X, et al. Murine model of elastase-induced proximal thoracic aortic aneurysm through a midline incision in the anterior neck. Front Cardiovasc Med. 2023;10:953514. doi:10.3389/fcvm.2023.953514

26. Richards DA, Aronovitz MJ, Calamaras TD, et al. Distinct Phenotypes Induced by Three Degrees of Transverse Aortic Constriction in Mice. Sci Rep. 2019;9:5844. doi:10.1038/s41598-019-42209-7

27. Pereira RO, Wende AR, Crum A, et al. Maintaining PGC-1α expression following pressure overload-induced cardiac hypertrophy preserves angiogenesis but not contractile or mitochondrial function. FASEB J. 2014;28(8):3691–3702. doi:10.1096/fj.14-253823

28. Riehle C, Wende AR, Zaha VG, et al. PGC-1β Deficiency Accelerates the Transition to Heart Failure in Pressure Overload Hypertrophy. Circ Res. 2011;109(7):783–793. doi:10.1161/CIRCRESAHA.111.243964

29. Rockman HA, Ross RS, Harris AN, et al. Segregation of atrial-specific and inducible expression of an atrial natriuretic factor transgene in an in vivo murine model of cardiac hypertrophy. Proc Natl Acad Sci U S A. 1991;88(18):8277–8281.

30. López B, Ravassa S, Moreno MU, et al. Diffuse myocardial fibrosis: mechanisms, diagnosis and therapeutic approaches. Nat Rev Cardiol. 2021;18(7):479–498. doi:10.1038/s41569-020-00504-1

31. Burkhoff D, Topkara VK, Sayer G, Uriel N. Reverse Remodeling With Left Ventricular Assist Devices. Circ Res. 2021;128(10):1594–1612. doi:10.1161/CIRCRESAHA.121.318160

32. Rodrigues PG, Miranda-Silva D, Li X, et al. The Degree of Cardiac Remodelling before Overload Relief Triggers Different Transcriptome and miRome Signatures during Reverse Remodelling (RR)—Molecular Signature Differ with the Extent of RR. Int J Mol Sci. 2020;21(24):9687. doi:10.3390/ijms21249687

33. Kassner A, Oezpeker C, Gummert J, et al. Mechanical circulatory support does not reduce advanced myocardial fibrosis in patients with end-stage heart failure. Eur J Heart Fail. 2021;23(2):324–334. doi:10.1002/ejhf.2021

34. Davidson LJ, Davidson CJ. Transcatheter Treatment of Valvular Heart Disease: A Review. JAMA. 2021;325(24):2480–2494. doi:10.1001/jama.2021.2133

35. Gao E, Lei YH, Shang X, et al. A Novel and Efficient Model of Coronary Artery Ligation and Myocardial Infarction in the Mouse. Circ Res. 2010;107(12):1445–1453. doi:10.1161/CIRCRESAHA.110.223925

36. Venema CS, van Bergeijk KeesH, Hadjicharalambous D, et al. Prediction of the Individual Aortic Stenosis Progression Rate and its Association With Clinical Outcomes. JACC Adv. 2024;3(4):100879. doi:10.1016/j.jacadv.2024.100879

37. Kim DH, Afilalo J, Shi SM, et al. Evaluation of Changes in Functional Status in the Year After Aortic Valve Replacement. JAMA Intern Med. 2019;179(3):383–391. doi:10.1001/jamainternmed.2018.6738

38. Schueler R, Sinning JM, Momcilovic D, et al. Three-Dimensional Speckle-Tracking Analysis of Left Ventricular Function after Transcatheter Aortic Valve Implantation. J Am Soc Echocardiogr. 2012;25(8):827–834.e1. doi:10.1016/j.echo.2012.04.023

39. Kim CA, Rasania SP, Afilalo J, Popma JJ, Lipsitz LA, Kim DH. Functional Status and Quality of Life After Transcatheter Aortic Valve Replacement. Ann Intern Med. 2014;160(4):243–254. doi:10.7326/M13-1316

40. Leon MB, Smith CR, Mack M, et al. Transcatheter Aortic-Valve Implantation for Aortic Stenosis in Patients Who Cannot Undergo Surgery. N Engl J Med. 2010;363(17):1597–1607. doi:10.1056/NEJMoa1008232

