## Supplemental figures for "Simplified and refined murine model of reversible aortic constriction for characterizing cardiac functional recovery"

**Supplemental Table 1 Cardiac function, geometry, and strain measured.** Aortic velocity, ejection fraction (EF), stroke volume (SV), cardiac output (CO) measured from 2D ultrasound. Wall thickness, left ventricle (LV) diameter and mass, and strain measured from 4D ultrasound. *$p$<0.05 compared to baseline, †$p$<0.05 compared to 3W TAC. Normally distributed data was analyzed using repeated measures ANOVA with Tukey's post hoc test, while non-parametric equivalents were used for non-normally distributed data. n=8/group.

| | | Baseline (Week 0) | 3W TAC (Week 3) | 1W deTAC (Week 4) | 4W deTAC (Week 7) |
|---|---|---|---|---|---|
| **Verification** | **Aortic Velocity** | 980 ± 121 | **3738 ± 396*** | **1781 ± 444*** | **1640 ± 312†** |
| **Function** | **EF (%)** | 62.0 ± 3.2 | **35.0 ± 7.9*** | **47.5 ± 8.8*†** | **58.8 ± 8.6†** |
| | **SV (μL)** | 28.3 ± 4.0 | **19.9 ± 4.5*** | **27.6 ± 5.1†** | **32.8 ± 6.5†** |
| | **CO (μL/min)** | 14.1 ± 2.0 | 11.0 ± 2.6 | 13.8 ± 2.7 | **17.5 ± 4.4†** |
| **Geometry** | **Wall Thickness (mm)** | 0.8 ± 0.2 | **1.1 ± 0.10*** | 0.9 ± 0.2 | 0.9 ± 0.1 |
| | **LV diameter (mm)** | 3.6 ± 0.4 | 4.1 ± 0.5 | 4.1 ± 0.5 | 4.1 ± 0.4 |
| | **LV mass (mg)** | 66.2 ± 7.9 | **94.4 ± 7.5*** | **85.8 ± 10.4*†** | **90.7 ± 7.6*** |
| | **LVM : BM (x10$^3$)** | 3.0 ± 0.4 | **4.0 ± 0.3*** | **3.5 ± 0.3†** | 3.6 ± 0.3 |
| **Peak Global Strain (%)** | **Circumferential ($E_{cc}$)** | -28.5 ± 3.5 | **-17.7 ± 4.0*** | **-23.4 ± 5.2†** | **-25.5 ± 4.0†** |
| | **Longitudinal ($E_{ll}$)** | -19.9 ± 3.0 | **-11.4 ± 3.3*** | -16.8 ± 3.4 | **-17.1 ± 2.0†** |
| | **Radial ($E_{rr}$)** | 24.3 ± 9.1 | **12.6 ± 4.5*** | **18.0 ± 3.2†** | 16.4 ± 5.2 |
| | **Surface Area ($E_{sa}$)** | -47.2 + 5.4 | **-28.8 + 6.8*** | **-40.0 + 8.2†** | **-41.8 + 5.5†** |

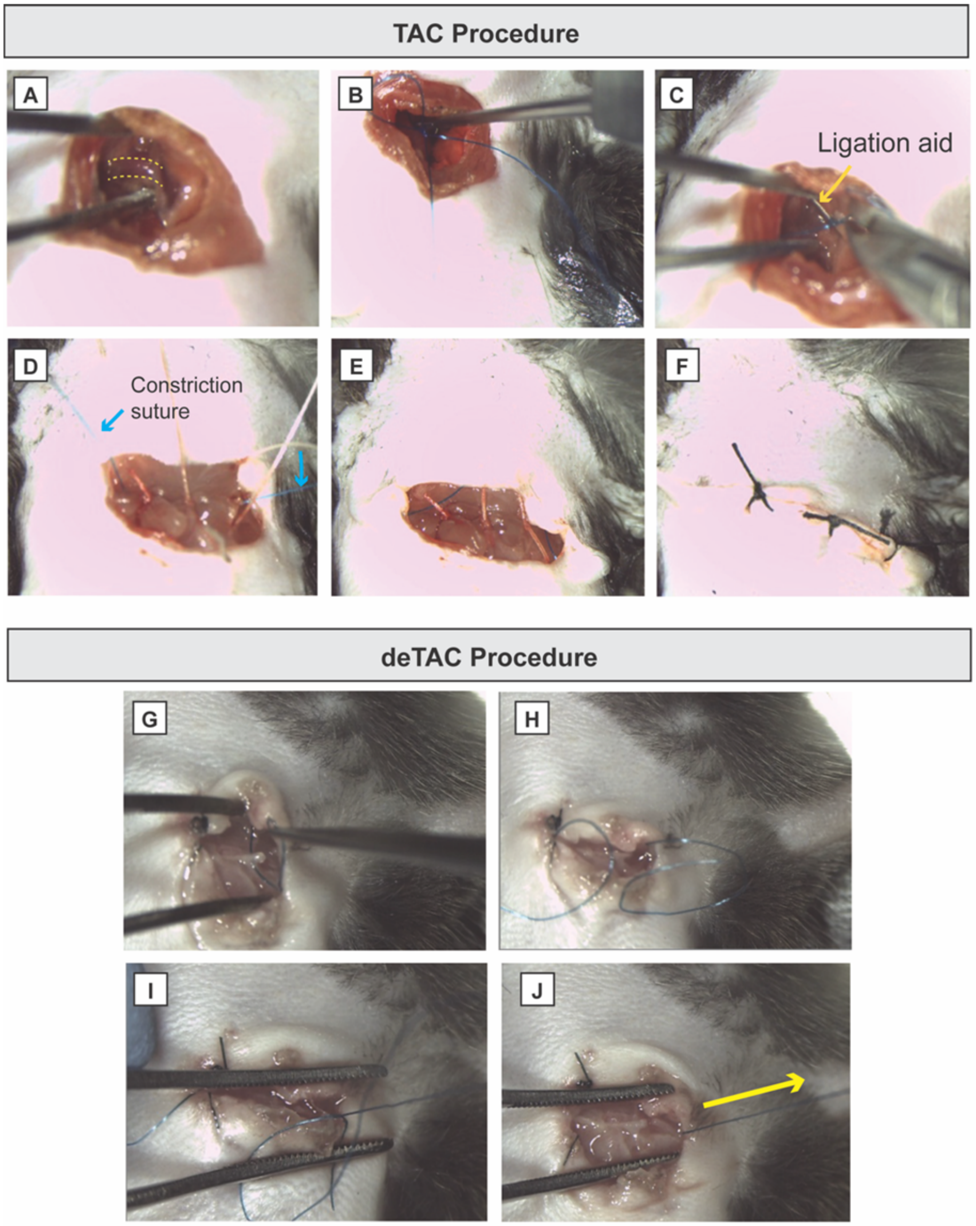


**S1. Surgical procedure for transverse aortic constriction (TAC) and aortic de-banding (deTAC).** TAC: **A)** Identify transverse aorta, **B-C)** Tie suture around aorta, **D-E)** close muscle layer with constriction suture accessible, **F)** close skin layer. deTAC: **G)** Open skin layer, **H-I)** access constriction suture, **J)** pull suture to release.

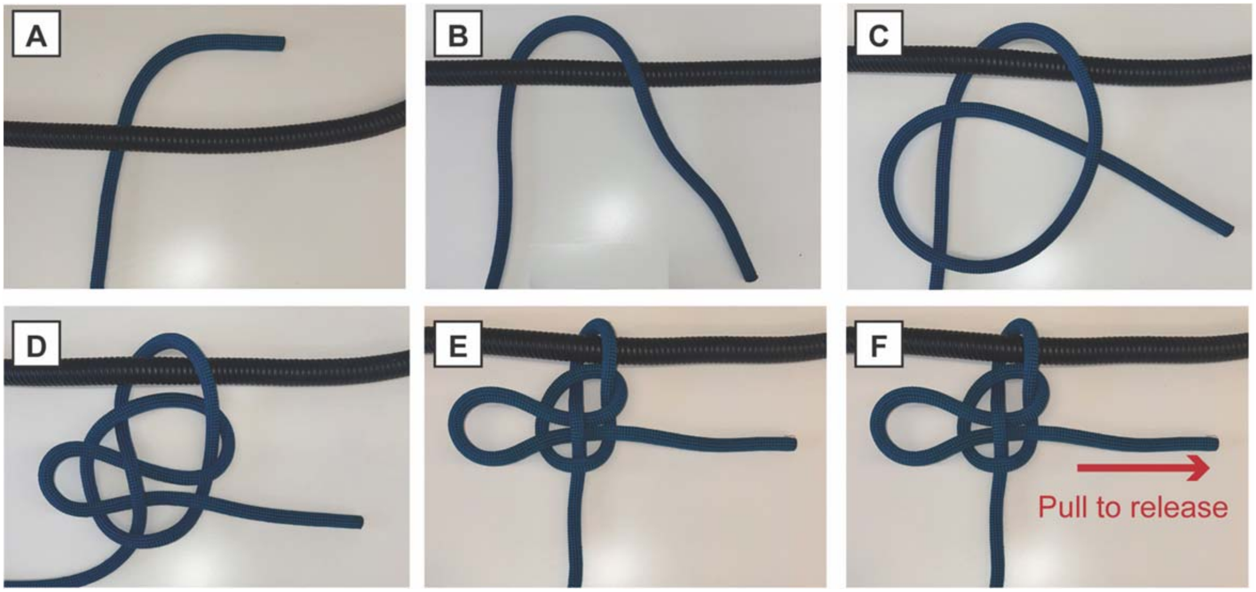

**S2. Schematic of constriction suture (mooring-hitch). A-E** Steps for aortic constriction. **F**. Final step enabling quick-release/deTAC and suture removal.

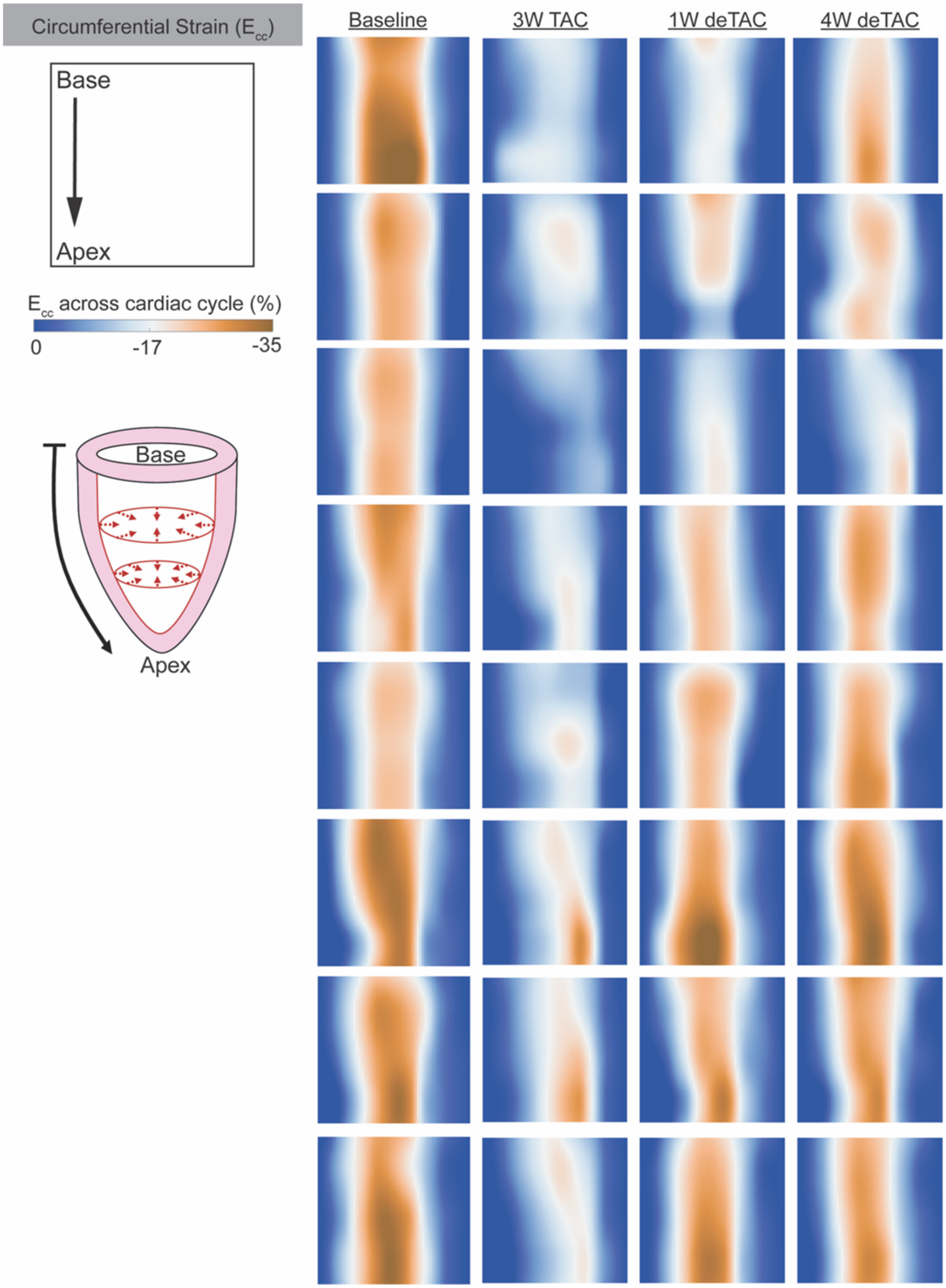


**S3. Individual Circumferential Strain Maps**

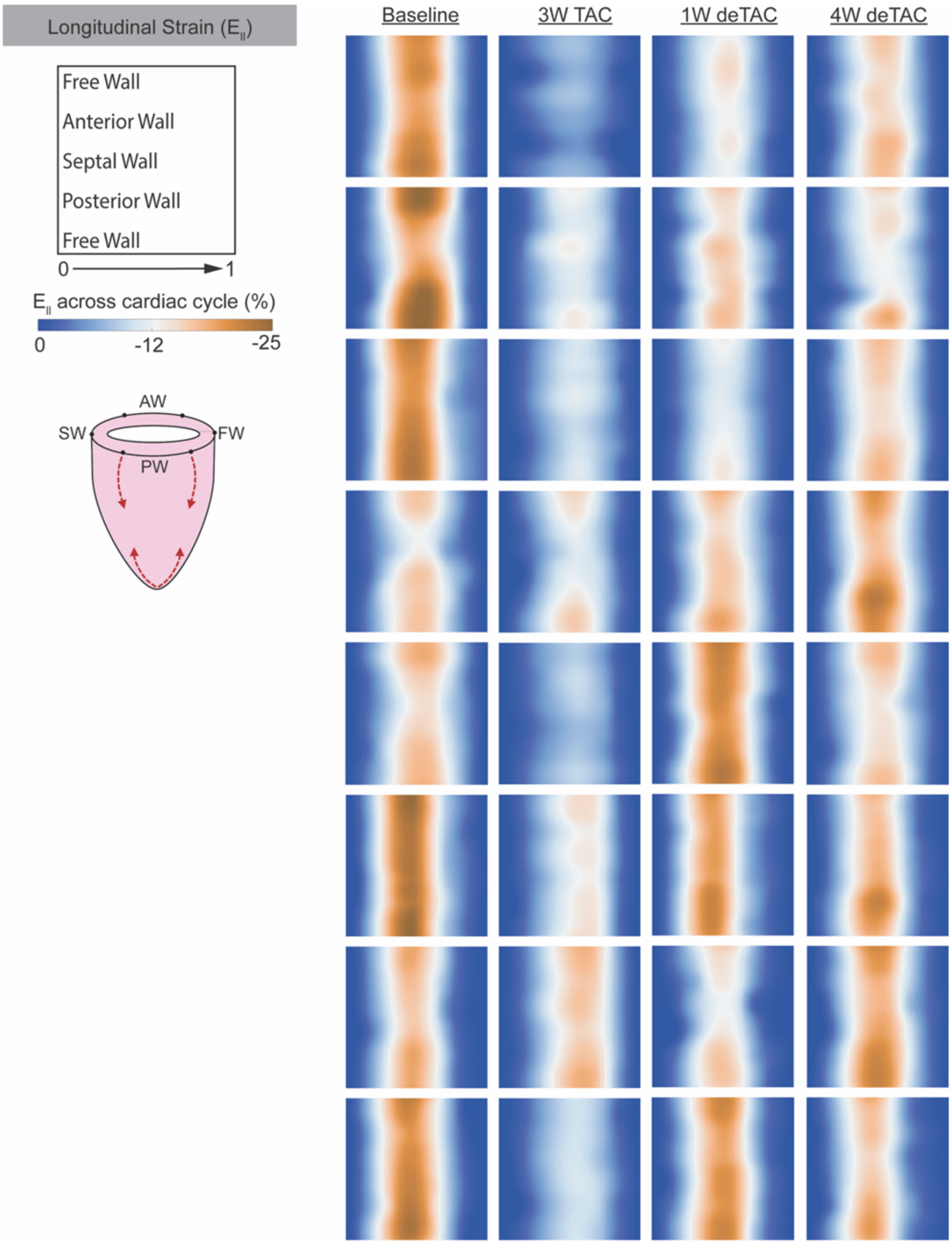


**S4. Individual Longitudinal Strain Maps**

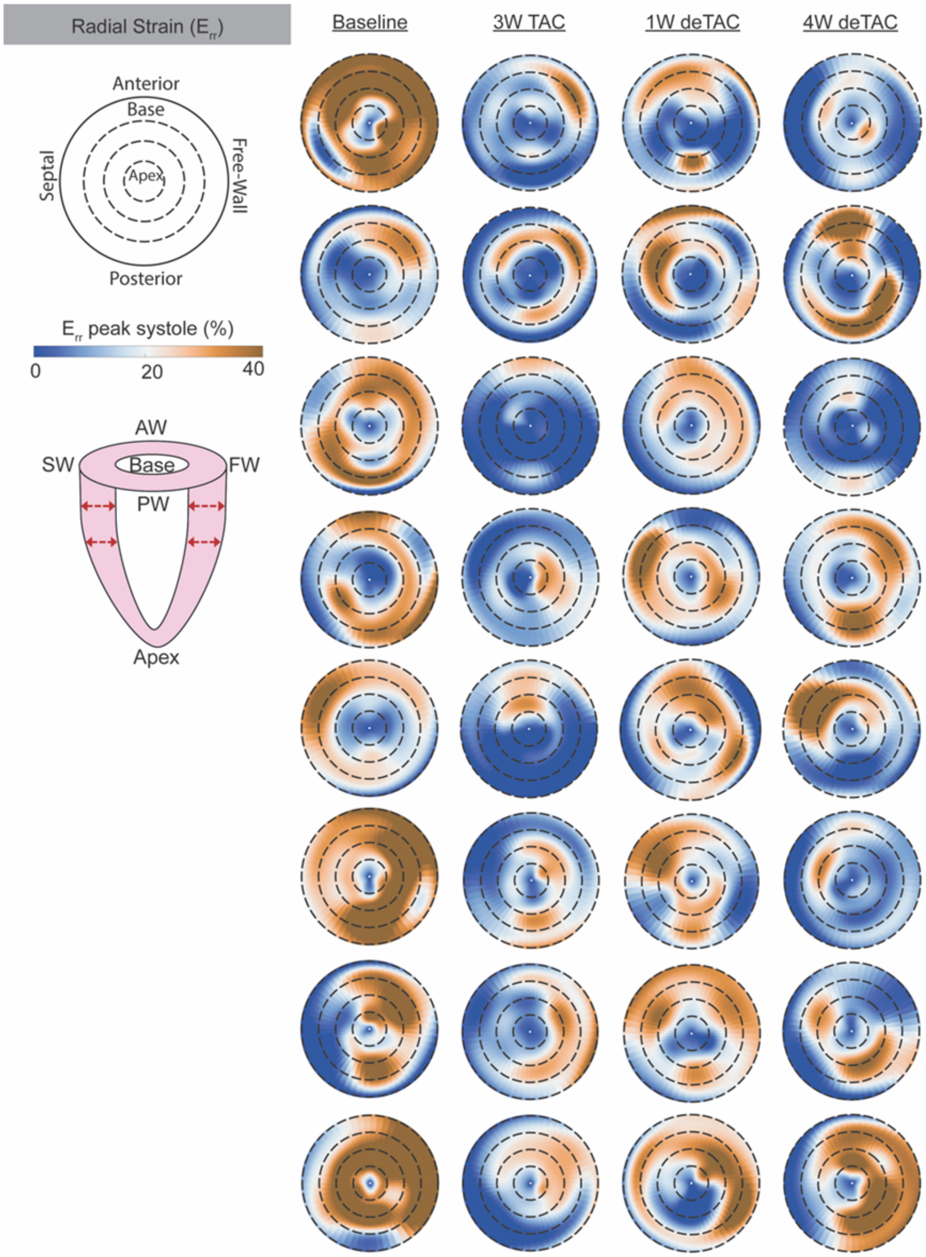


**S5. Individual Radial Strain Maps**

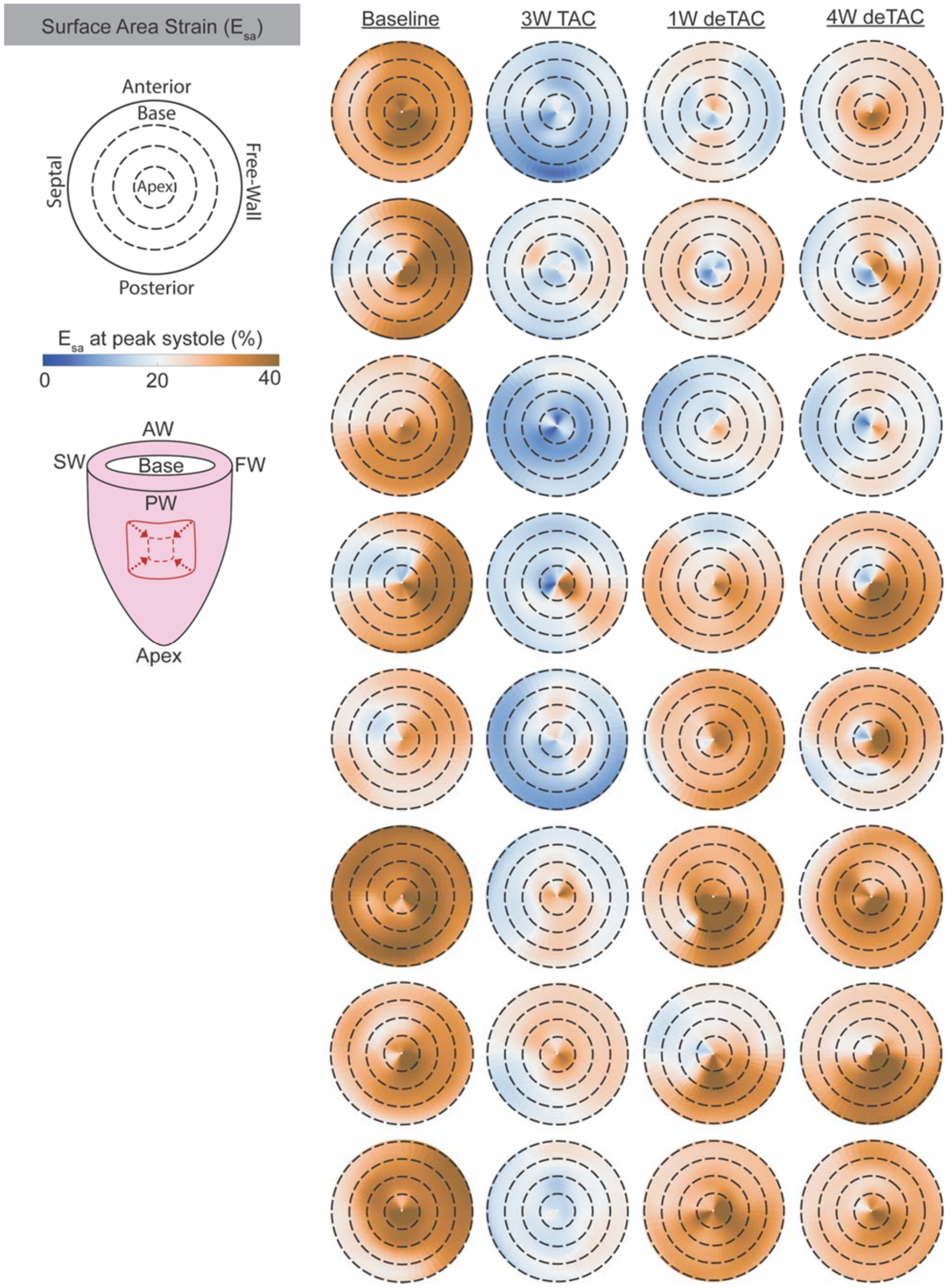


**S6. Individual Surface Area Strain Maps**
